# Impact of dietary synbiotics on growth performance, gut morphology, and immune function, and welfare indicators of broiler chickens with or without early life antibiotic supplementation via drinking water

**DOI:** 10.1101/2025.10.30.685693

**Authors:** M.J.M. Musny, K.K.K.P. Perera, H.E.L. De Seram, P. Weththasinghe, N.D. Karunaratne

**Author notes:** These authors contributed equally to this work (M.J.M. Musny, K.K.K.P. Perera).

## Abstract

The widespread use of antibiotics in broiler production has contributed to antimicrobial resistance, necessitating the development of sustainable alternatives. Synbiotics, combining probiotics and prebiotics in a synergistic formulation, have emerged as promising candidates to replace antibiotics. This study evaluated the efficacy of dietary synbiotics as alternatives to early-life antibiotic supplementation in broiler chickens, examining their effects on growth performance, gut morphology, and welfare. A total of 80 mixed-sex Cobb 500 broiler chickens were allocated to a 2 × 2 factorial design investigating synbiotic supplementation (*Bacillus subtilis* and dried fermented *Saccharomyces cerevisiae* extract) and enrofloxacin treatment (days 0-5) administered via drinking water. During the initial growth period (days 0-7), synbiotic supplementation reduced body weight gain compared to controls. However, synbiotic × antibiotic interactions were observed during days 7-14 and 14-21, where synbiotics without antibiotics produced the highest weight gain in the second week, while controls without either treatment achieved superior performance in the third week. Feed intake was increased by synbiotic supplementation during days 7-14, while antibiotic treatment consistently elevated feed consumption throughout multiple periods and overall trial duration. Feed conversion efficiency was initially impaired by synbiotics (days 0-7) but improved during days 14-21. Antibiotic supplementation resulted in a poorer overall feed conversion ratio. Gut morphological analysis revealed significant synbiotic × antibiotic interactions for duodenal length and empty weights of duodenum and caeca, with synbiotics enhancing these parameters only when combined with antibiotic treatment. Both synbiotic and antibiotic supplementation independently reduced ileal content weight, while antibiotic treatment specifically decreased ileal weight and caecal length. Immune function evaluation through heterophil-to-lymphocyte ratios demonstrated a tendency for interaction, where synbiotics reduced this ratio in the absence of antibiotics but showed no effect when combined with antibiotic treatment. Neither treatment affected spleen or bursa weights, indicating minimal impact on immune organ development. These findings suggest that while dietary synbiotics show promise as alternatives to antibiotics in broiler production, their efficacy varies depending on concurrent antibiotic exposure and growth phase. Synbiotics demonstrated beneficial effects on gut morphology and stress markers, particularly when administered without concurrent antibiotic treatment, supporting their potential as functional alternatives in antimicrobial-free poultry production.

## Introduction

Global food security and the nutritional demands of an expanding world population represent critical challenges in contemporary agriculture (Hussain et al., 2025). With per capita protein consumption rising globally and poultry products constituting approximately 70% of total animal protein intake, the development of sustainable production systems has become imperative. Poultry production has gained prominence as an efficient means of converting feed resources into high-quality animal protein, establishing itself as a cornerstone of global food security strategies (Ayalew et al., 2022). Broiler production has experienced exponential growth globally due to its comparative advantages, including high nutritional quality, appealing taste, low fat content, short production cycle, low production cost, and affordability even for lower-income populations (Ayalew et al., 2022). In Sri Lanka, this trend is evident with steady growth in local poultry production (Livestock Statistical Bulletin, 2024).

Traditionally, antibiotics have been used extensively in the poultry industry for decades to control infectious diseases while promoting broiler growth. Supplementing broiler diets with antibiotics through feed has been shown to increase body weight gain by 5.8%, mainly through regulation of intestinal microflora and improved feed conversion efficiency (Ayalew et al., 2022; Mojarla et al., 2025). However, the widespread use of in-feed antibiotics in broiler farming has led to the rise of antimicrobial-resistant bacteria, posing a serious public health threat (Chen et al., 2025). Antimicrobial resistance has become a major global health concern directly linked to the overuse and misuse of antibiotics in poultry farming. Additionally, antibiotic residues in animal products and environmental contamination have raised significant concerns regarding human food safety (Ayalew et al., 2022). Consequently, the use of antibiotic growth promoters has been banned in poultry feed worldwide, creating an urgent need to identify suitable alternatives to antibiotics in modern poultry farming (Ayalew et al., 2022; Mojarla et al., 2025).

As alternatives to antibiotics, synbiotics have emerged as a promising solution (Gadde et al., 2017). Synbiotics are a synergistic combination of probiotics and prebiotics designed to promote host health and well-being by enhancing the survival, colonization, and function of probiotics in the gastrointestinal tract while supporting the host’s natural gut microbiota (Mojarla et al., 2025). Probiotics are live microorganisms that provide health benefits when administered in adequate amounts, while prebiotics are non-digestible food components that selectively stimulate the growth and activity of beneficial gut bacteria. Microbe species belonging to *Lactobacillus, Bacillus, Streptococcus, Enterococcus, Bifidobacterium, Aspergillus,* and *Saccharomyces* are commonly used as probiotics in the poultry industry (Olmez et al., 2025). The most popular prebiotics include oligosaccharides such as galactose-oligosaccharides, fructose-oligosaccharides, mannan-oligosaccharides, raffinose, lactulose, inulin, stachyose, and chitosan-oligosaccharides (Olmez et al., 2025). Additionally, the wall of *Saccharomyces cerevisiae* can also be used as a prebiotic due to its content, including α-mannan, β-glucan, and chitin (Al-Baadani et al., 2025).

Extensive research has established synbiotics as efficacious alternatives to antimicrobial growth promoters, demonstrating significant improvements in growth performance, intestinal morphology, beneficial microbiota proliferation, and immune function in broiler chickens (Mojarla et al., 2025; Olmez et al., 2025; Song et al., 2022). These bioactive compounds enhance zootechnical performance and feed utilization efficiency through modulation of various physiological pathways, although the precise mechanisms underlying these effects remain incompletely elucidated (Reuben et al., 2021). Furthermore, synbiotic supplementation has demonstrated ameliorative effects on stress-related parameters in broilers subjected to thermal stress (Hu et al., 2022) and elevated stocking densities (Kridtayopas et al., 2019; Ragab et al., 2024). Contemporary broiler genotypes, selected for accelerated growth rates and enhanced carcass characteristics, are commonly reared under intensive management systems characterized by high stocking densities, conditions that may compromise overall welfare status (Meluzzi & Sirri, 2009).

Despite the documented benefits of dietary synbiotics, the interactive effects of synbiotic supplementation in conjunction with or independent of early-life antimicrobial administration remain poorly characterized, particularly within the Sri Lankan production context. A significant knowledge gap exists in the literature regarding the comparative efficacy of synbiotics as functional replacements for antibiotics, specifically concerning broiler performance parameters, nutrient digestibility, immunomodulatory responses, and welfare indices. Contemporary research continues to investigate optimal administration protocols and formulations to maximize the therapeutic potential of these alternatives, establishing their viability for antimicrobial-free poultry production systems.

This study aims to evaluate the effects of dietary synbiotics as a potential alternative to antibiotics administered in drinking water during the first five days of life in broiler chickens, specifically examining their impact on growth performance, gut morphology, immune function, and stress indicators. This study hypothesizes that dietary synbiotics will improve growth performance, gut morphology, and immune function while reducing stress in broiler chickens with or without early-life antibiotic supplementation via drinking water. The findings of this study will be important for improving food safety, supporting sustainable poultry farming, and promoting responsible use of antibiotics in the poultry industry in Sri Lanka and worldwide.

## Materials and methods

### Ethical approval

Ethical approval for the experimental procedure was obtained from the Committee for the Ethical Clearance on Animal Research of the Faculty of Veterinary Medicine and Animal Science, University of Peradeniya, Sri Lanka (Ethical clearance certificate number: VERC_25_12).

### Birds and housing

A total of 80 mixed-sex Cobb 500 broiler chickens were procured from a commercial hatchery and housed in battery cages measuring 31.5 cm × 31.0 cm × 19.5 cm (length × width × height). The cages were equipped with wire mesh flooring with a grid size of 0.8 cm × 0.8 cm and arranged in two rows with two levels each, within an open-sided housing facility.

Environmental conditions were carefully controlled throughout the experimental period. The ambient temperature was initially set at 32°C on day 0 and subsequently reduced gradually based on behavioural observations of the birds. Photoperiod was maintained at 24 hours of continuous lighting from day 0 to 7, followed by a 12-hour light cycle from day 8 to 35. Light intensity was sustained at a minimum of 25 lux throughout the trial period. Feed and water were provided *ad libitum* for the duration of the study.

Feeding and watering systems were adapted according to the developmental stage of the birds. During the initial brooding period (day 0-7), each cage was equipped with a tray feeder (8.8 cm diameter) and a mini bell drinker (1 L capacity). From day 8 to 35, these were replaced with turbo feeders (10.2 cm pan diameter) and manual bell drinkers (2 L capacity). Each cage was additionally fitted with a 75 W electric bulb positioned at the cage top level to provide supplemental lighting and brooding heat. To maintain optimal brooding temperature during the first two weeks, room windows were covered with polyethylene sheeting.

The experimental design employed a randomized complete block design with four replications per treatment, housing five birds per cage. On day 0, birds were weighed on a cage basis and allocated to maintain uniform initial body weight distribution across treatments. Initial feed allocation to each cage was measured and recorded at the commencement of the trial.

### Experimental treatments

The experimental treatments were arranged in a 2 × 2 factorial design, with two main factors: dietary synbiotic supplementation (presence or absence) and antibiotic administration via drinking water during days 0-5 (presence or absence). The synbiotic supplement utilized was Pro Bio XP 100 (Bio Nutri International Pvt Ltd, Moratuwa, Sri Lanka), containing *Bacillus subtilis* as the probiotic component and dried, fermented *Saccharomyces cerevisiae* extract as the prebiotic component. The antibiotic treatment consisted of Enrofloxacin, a fluoroquinolone-class water-soluble powder formulated for poultry use.

All experimental diets were formulated to meet or exceed the nutritional requirements specified for Cobb 500 broilers (Cobb-Vantress, 2022) and were formulated to be iso-energetic and iso-nitrogenous across treatments. A three-phase feeding program was implemented: broiler starter feed (days 0-10) provided in crumble form, broiler grower feed (days 11-21) in mash form, and broiler finisher feed (days 22-35) in mash form. The complete ingredient composition and calculated nutrient profiles are detailed in Table 1. The synbiotic supplement was incorporated into the designated experimental feeds during the manufacturing process to ensure homogeneous distribution throughout the diet.

**Table 1.** Feed ingredient composition and calculated nutrient levels of the experimental diets (as-is basis)

| **Ingredient (%)** | **Starter** | **Grower** | **Finisher** |
| --- | --- | --- | --- |
| **Maize** | 51.35 | 54.45 | 57.75 |
| **Soybean meal** | 34.40 | 29.50 | 24.50 |
| **Fish meal** | 5.00 | 4.00 | 3.00 |
| **Palm oil** | 3.00 | 4.00 | 6.00 |
| **Rice bran** | 1.00 | 3.00 | 4.00 |
| **Lysine** | 0.70 | 0.70 | 0.55 |
| **Methionine** | 0.40 | 0.40 | 0.40 |
| **Threonine** | 0.10 | 0.10 | 0.05 |
| **Limestone** | 1.00 | 1.00 | 1.00 |
| **Dicalcium phosphate** | 1.80 | 1.60 | 1.50 |
| **Salt** | 0.35 | 0.35 | 0.35 |
| **Vitamin-mineral premix** | 0.50 | 0.50 | 0.50 |
| **Choline chloride** | 0.10 | 0.10 | 0.10 |
| **Coccidiostat** | 0.05 | 0.05 | 0.05 |
| **Enzyme cocktail** | 0.05 | 0.05 | 0.05 |
| **Toxin binder** | 0.10 | 0.10 | 0.10 |
| **Pro-bio (synbiotic)** | 0.10 | 0.10 | 0.10 |
| **<u>Calculated nutrient levels (%)</u>** |  |  |  |
| **Metabolizable energy (kcal/kg)** | 3,050 | 3,150 | 3,280 |
| **Crude protein** | 22.8 | 20.5 | 18.2 |
| **Lysine** | 1.40 | 1.25 | 1.05 |
| **Methionine** | 0.68 | 0.66 | 0.63 |
| **Methionine + Cystine** | 1.05 | 0.96 | 0.88 |
| **Threonine** | 0.86 | 0.78 | 0.68 |
| **Tryptophan** | 0.23 | 0.20 | 0.18 |
| **Calcium** | 1.02 | 0.96 | 0.90 |
| **Available phosphorus** | 0.48 | 0.45 | 0.42 |
| **Sodium** | 0.18 | 0.17 | 0.16 |
| **Chloride** | 0.23 | 0.22 | 0.22 |
| **Linoleic Acid** | 1.65 | 1.85 | 2.20 |

### Data and sample collection

#### Performance parameters

Body weight and feed intake were monitored on a cage basis at 7-day intervals (days 7, 14, 21, 28, and 35 post-hatch). Body weight gain and feed conversion ratio were subsequently calculated from these measurements. Daily mortality records were maintained, with comprehensive postmortem examinations conducted on all deceased birds.

#### Blood sample collection

On day 35, two birds were randomly selected from each cage, and blood samples (0.2-0.3 mL) were collected from the brachial vein using a 3 mL syringe fitted with a 23-gauge needle.

Prior to venipuncture, birds were properly restrained, and the sampling site was disinfected with 70% isopropyl alcohol. Blood smears were prepared immediately upon collection, labelled appropriately, and air-dried for subsequent analysis.

#### Gastro-intestinal tract morphology

Two birds per cage were randomly selected and humanely euthanized by jugular vein severance following electrical stunning. Individual body weights were recorded immediately before euthanization. The complete digestive tract was systematically removed and dissected into constituent components, including proventriculus, gizzard, duodenum, jejunum, ileum, and caeca. Associated organs, including the pancreas, liver, spleen, and bursa of Fabricius, were also excised and weighed.

For the duodenum, jejunum, ileum, and caeca, both full weight (with contents) and empty weight measurements were recorded, along with length measurements. Full weights were documented for the proventriculus and gizzard, while absolute weights were recorded for the pancreas, liver, spleen, and bursa of Fabricius. Content weights for each intestinal segment were determined by calculating the difference between full and empty weights.

Composite small intestine parameters (full weight, empty weight, length, and content weight) were derived by summing the corresponding measurements from the duodenum, jejunum, and ileum. All organ weights and length measurements were normalized to individual body weight to obtain relative values expressed as proportions of total body weight, enabling standardized morphometric comparisons.

### Laboratory procedures

#### Proximate analysis of feed

Starter, grower, and finisher feeds (formulated without synbiotic supplementation) utilized in the experimental protocol were subjected to proximate composition analysis. Moisture content was determined following the standardized methodology outlined by the Association of Official Analytical Chemists (AOAC, 1995, method 930.15). Nitrogen content was quantified according to the AOAC method 990.03 (1995), with crude protein calculated using a nitrogen-to-protein conversion factor of 6.25. Ether extract (crude fat) was analyzed following the AOAC method 920.39 (1995). Ash content was determined according to the AOAC method 942.05 (1995), and crude fiber content was quantified using the AOAC method 991.43 (1995).

#### Heterophil-to-lymphocyte ratio assessment

The air-dried blood smears were stained using Leishman’s stain (L6254-25G, Sigma-Aldrich, Merck Life Science Pvt Ltd, VIC, Australia) according to standard hematological procedures. Microscopic examination was conducted using a light microscope (Olympus CX23, Hubei, China) equipped with oil immersion at ×100 magnification. For each blood smear, differential leukocyte counts were performed by systematically examining and enumerating 200 white blood cells per slide. Within this population, heterophils and lymphocytes were identified based on morphological characteristics and quantified. The heterophil-to-lymphocyte ratio (H:L ratio) was calculated as a stress indicator by dividing the number of heterophils by the number of lymphocytes and expressing the result as a percentage, following the methodology described by Gross and Siegel (1983).

### Data analysis

The experiment was a randomized complete block design with a cage level used as a block to account for potential environmental differences between the 2 levels. Data were analyzed using STATA statistical software, and 2-way analysis of variance was used to determine the main effects of, and interaction between, synbiotic and antibiotic supplementation (StataCorp LLC, College Station, TX). The significance level was *P* ≤ 0.05, and trends were considered when 0.10 ≥ *P* > 0.05. Mean separation was completed using the Tukey–Kramer test. Data were tested for normality using the Shapiro–Wilk test before analyzing variance, and the Mann-Whitney test was used when data were not normally distributed.

## Results

The analyzed nutrient composition of the starter, grower and finisher feed (without synbiotics) used for the experiment is presented in Table 2.

**Table 2.** Analyzed nutrient composition of the experimental diets (as-is basis)

| **Nutrient (%)** | **Starter** | **Grower** | **Finisher** |
| --- | --- | --- | --- |
| **Crude protein** | 23.4 | 21.6 | 17.6 |
| **Ether extract** | 3.4 | 7.2 | 8.2 |
| **Crude fiber** | 6.3 | 6.5 | 6.5 |
| **Ash** | 8.0 | 7.3 | 5.6 |
| **Moisture** | 14.9 | 11.0 | 10.1 |

The effects of dietary synbiotics and early-life antibiotics supplementation on body weight gain of broiler chickens are presented in Table 3. A significant main effect of synbiotics was observed during the initial growth period (Days 0-7). During days 0-7, chickens receiving synbiotics had lower body weight gain compared to those without synbiotics (P = 0.056). A significant synbiotic and antibiotic interaction was observed during days 7-14 (P = 0.010) and a trend during days 14-21 (P = 0.086). During days 7-14, the combination of synbiotics without antibiotics produced the highest weight gain, while the control group showed the lowest weight gain. The antibiotic treatment groups showed intermediate performance regardless of synbiotic supplementation. During days 14-21, chickens receiving neither synbiotics nor antibiotics achieved the highest weight gain, while those receiving synbiotics without antibiotics had the lowest gain. No significant main effects or interactions were observed for the later growth periods or overall performance.

**Table 3.** Effects of dietary synbiotics and early-life antibiotics supplementation on body weight gain of broiler chickens.

| Synbiotic | Antibiotic | Body weight gain (g) |  |  |  |  |  |
| --- | --- | --- | --- | --- | --- | --- | --- |
|  |  | Day 0-7 | Day 7-14 | Day 14-21 | Day 21-28 | Day 28-35 | Day 0-35 |
| No | No | 123.7 | 177.7<sup>c</sup> | 494.9<sup>a</sup> | 568.6 | 642.9 | 2007.7 |
|  | Yes | 121.8 | 199.4<sup>bc</sup> | 474.0<sup>ab</sup> | 574.8 | 670.2 | 2040.2 |
| Yes | No | 111.6 | 253.1<sup>a</sup> | 417.6<sup>b</sup> | 575.3 | 687.2 | 2044.7 |
|  | Yes | 111.0 | 201.9<sup>b</sup> | 455.7<sup>ab</sup> | 565.5 | 680.1 | 2014.3 |
| <sup>1</sup>SEM |  | 4.48 | 14.13 | 17.16 | 30.56 | 27.43 | 51.87 |
| **Main Effects** |  |  |  |  |  |  |  |
| **<u>Synbiotic</u>** |  |  |  |  |  |  |  |
| No |  | 122.7<sup>a</sup> | 188.6 | 484.5 | 571.7 | 656.5 | 2023.9 |
| Yes |  | 111.3<sup>b</sup> | 227.5 | 436.7 | 570.4 | 683.6 | 2029.5 |
| **<u>Antibiotic</u>** |  |  |  |  |  |  |  |
| No |  | 117.6 | 215.4 | 456.3 | 571.9 | 665.0 | 2026.2 |
| Yes |  | 116.4 | 200.7 | 464.9 | 570.2 | 675.2 | 2027.2 |
| **<u>Probability</u>** |  |  |  |  |  |  |  |
| Synbiotic |  | 0.056 | <0.001 | 0.001 | 0.877 | 0.254 | 0.614 |
| Antibiotic |  | 0.764 | 0.277 | 0.390 | 0.886 | 0.481 | 0.658 |
| Synbiotic × Antibiotic |  | 0.880 | 0.010 | 0.086 | 0.794 | 0.531 | 0.544 |

<sup>1</sup>SEM-Standard Error of Mean (n=4).
<sup>a-c</sup>Means within a main effect or interaction not sharing a common superscript are significantly different ( $P \leq 0.05$ ).

Feed intake patterns throughout the experimental period are shown in Table 4. Significant main effects of synbiotics were limited to the days 7-14 period, when synbiotic supplementation increased feed intake compared to the control (P = 0.008). Antibiotic supplementation demonstrated more consistent effects on feed intake. Significant increases in feed intake were observed during days 7-14 (P = 0.042), days 21-28 (P = 0.091), and for the overall period (P = 0.012) in antibiotic-supplemented birds compared to those without antibiotics. No significant synbiotic and antibiotic interactions were detected for feed intake at any time period.

**Table 4.** Effects of dietary synbiotics and early-life antibiotics supplementation on feed intake of broiler chickens.

| Synbiotic | Antibiotic | Feed intake (g) |  |  |  |  |  |
| --- | --- | --- | --- | --- | --- | --- | --- |
|  |  | Day 0-7 | Day 7-14 | Day 14-21 | Day 21-28 | Day 28-35 | Day 0-35 |
| No | No | 286.1 | 256.8 | 649.1 | 824.7 | 1013.1 | 3029.7 |
|  | Yes | 298.6 | 334.7 | 642.7 | 979.9 | 1087.2 | 3343.0 |
| Yes | No | 290.9 | 359.1 | 704.7 | 829.5 | 1044.2 | 3228.3 |
|  | Yes | 268.0 | 381.9 | 694.4 | 839.8 | 1094.3 | 3278.4 |
| <sup>1</sup>SEM |  | 15.84 | 29.91 | 36.01 | 64.95 | 37.84 | 110.57 |
| **Main Effects** |  |  |  |  |  |  |  |
| **Synbiotic** |  |  |  |  |  |  |  |
| No |  | 292.3 | 295.7<sup>b</sup> | 645.9 | 902.3 | 1050.1 | 3186.3 |
| Yes |  | 279.5 | 370.5<sup>a</sup> | 699.5 | 834.6 | 1069.3 | 3253.3 |
| **Antibiotic** |  |  |  |  |  |  |  |
| No |  | 288.5 | 307.9<sup>b</sup> | 676.9 | 827.1<sup>b</sup> | 1028.6 | 3129.0<sup>b</sup> |
| Yes |  | 283.3 | 358.3<sup>a</sup> | 668.5 | 909.8<sup>a</sup> | 1090.7 | 3310.7<sup>a</sup> |
| **Probability** |  |  |  |  |  |  |  |
| Synbiotic |  | 0.819 | 0.008 | 0.275 | 0.959 | 0.529 | 0.112 |
| Antibiotic |  | 0.553 | 0.042 | 0.901 | 0.091 | 0.134 | 0.012 |
| Synbiotic × Antibiotic |  | 0.236 | 0.310 | 0.956 | 0.265 | 0.732 | 0.136 |

<sup>1</sup>SEM-Standard Error of Mean (n=4).
<sup>a-b</sup>Means within a main effect or interaction not sharing a common superscript are significantly different (P≤0.05).

Feed conversion ratio (FCR) data are presented in Table 5, showing the efficiency of feed utilization across different growth periods. Synbiotic supplementation significantly affected FCR during the early growth period (days 0-7), where synbiotic-supplemented birds had a poorer conversion ratio than controls (P = 0.086). This trend reversed during days 14-21, with synbiotic groups showing improved FCR versus control (P = 0.024). Antibiotic supplementation demonstrated a significant effect on overall feed conversion efficiency, where antibiotic-treated birds had a poorer FCR compared to those without antibiotics (P = 0.075).

**Table 5.**
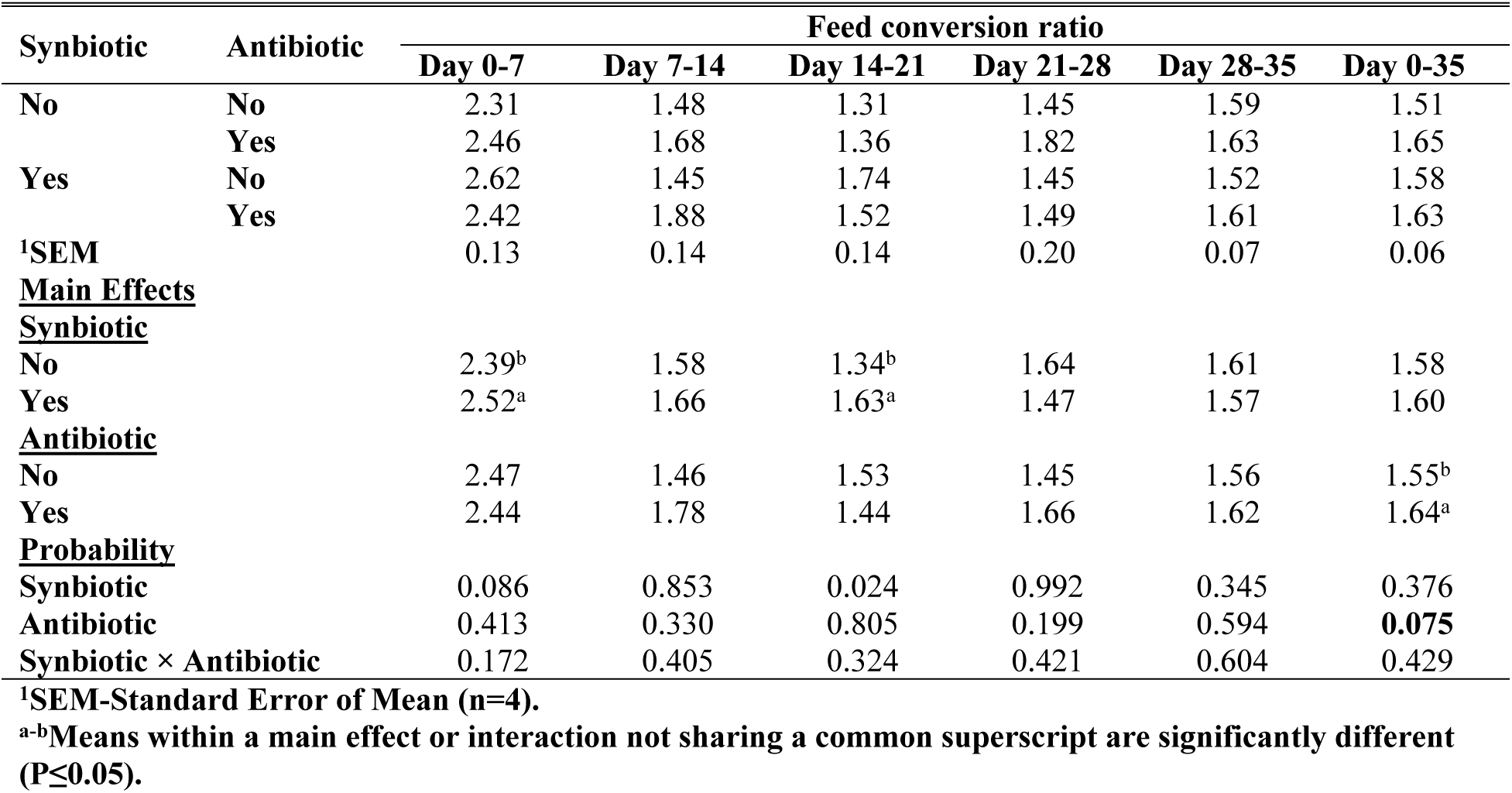
Effects of dietary synbiotics and early-life antibiotics supplementation on the feed conversion ratio of broiler chickens.

| Synbiotic | Antibiotic | Feed conversion ratio |  |  |  |  |  |
| --- | --- | --- | --- | --- | --- | --- | --- |
|  |  | Day 0-7 | Day 7-14 | Day 14-21 | Day 21-28 | Day 28-35 | Day 0-35 |
| No | No | 2.31 | 1.48 | 1.31 | 1.45 | 1.59 | 1.51 |
|  | Yes | 2.46 | 1.68 | 1.36 | 1.82 | 1.63 | 1.65 |
| Yes | No | 2.62 | 1.45 | 1.74 | 1.45 | 1.52 | 1.58 |
|  | Yes | 2.42 | 1.88 | 1.52 | 1.49 | 1.61 | 1.63 |
| <sup>1</sup>SEM |  | 0.13 | 0.14 | 0.14 | 0.20 | 0.07 | 0.06 |
| **Main Effects** |  |  |  |  |  |  |  |
| **<u>Synbiotic</u>** |  |  |  |  |  |  |  |
| No |  | 2.39<sup>b</sup> | 1.58 | 1.34<sup>b</sup> | 1.64 | 1.61 | 1.58 |
| Yes |  | 2.52<sup>a</sup> | 1.66 | 1.63<sup>a</sup> | 1.47 | 1.57 | 1.60 |
| **<u>Antibiotic</u>** |  |  |  |  |  |  |  |
| No |  | 2.47 | 1.46 | 1.53 | 1.45 | 1.56 | 1.55<sup>b</sup> |
| Yes |  | 2.44 | 1.78 | 1.44 | 1.66 | 1.62 | 1.64<sup>a</sup> |
| **<u>Probability</u>** |  |  |  |  |  |  |  |
| Synbiotic |  | 0.086 | 0.853 | 0.024 | 0.992 | 0.345 | 0.376 |
| Antibiotic |  | 0.413 | 0.330 | 0.805 | 0.199 | 0.594 | **0.075** |
| Synbiotic × Antibiotic |  | 0.172 | 0.405 | 0.324 | 0.421 | 0.604 | 0.429 |

<sup>1</sup>SEM-Standard Error of Mean (n=4).
<sup>a-b</sup>Means within a main effect or interaction not sharing a common superscript are significantly different (P≤0.05).

The effects of dietary synbiotics and early-life antibiotic supplementation on digestive tract weights relative to body weight are presented in Table 6. A significant synbiotic and antibiotic interaction was detected for empty duodenal weight (P = 0.030) and empty caecal weight (P = 0.035). In both cases, synbiotic supplementation increased the relative weights only when combined with antibiotic treatment, whereas no effect was observed in the absence of antibiotics. No other interactions were significant. Among the main effects, antibiotic supplementation reduced ileal weight (P = 0.017), while synbiotics did not significantly influence any digestive organ weights. The weights of the pancreas, proventriculus, gizzard, jejunum, small intestine, and other segments were unaffected by either treatment.

**Table 6.** Effects of dietary synbiotics and early-life antibiotics supplementation on digestive tract weights (proportional to body weight) of broiler chicken aged 36 days.

| Synbiotic | Antibiotic | Pan (%) | Full weights (%) |  |  |  |  |  |  | Empty weights (%) |  |  |  |  |
| --- | --- | --- | --- | --- | --- | --- | --- | --- | --- | --- | --- | --- | --- | --- |
|  |  |  | <sup>1</sup>Pro | Giz | Duo | Jej | Ile | Cae | SI | Duo | Jej | Ile | Cae | SI |
| No | No | 0.21 | 0.36 | 2.88 | 0.52 | 1.17 | 1.21 | 0.55 | 2.89 | 0.38<sup>ab</sup> | 0.77 | 0.75 | 0.30<sup>ab</sup> | 1.91 |
|  | Yes | 0.22 | 0.35 | 2.81 | 0.49 | 1.33 | 0.98 | 0.48 | 2.80 | 0.38<sup>ab</sup> | 0.84 | 0.73 | 0.28<sup>b</sup> | 1.95 |
| Yes | No | 0.19 | 0.34 | 2.71 | 0.48 | 1.15 | 1.03 | 0.49 | 2.66 | 0.33<sup>b</sup> | 0.75 | 0.72 | 0.30<sup>ab</sup> | 1.80 |
|  | Yes | 0.20 | 0.36 | 2.87 | 0.52 | 1.27 | 1.02 | 0.56 | 2.81 | 0.41<sup>a</sup> | 0.83 | 0.79 | 0.36<sup>a</sup> | 2.03 |
| <sup>2</sup>SEM |  | 0.014 | 0.024 | 0.249 | 0.496 | 1.212 | 1.028 | 0.523 | 2.736 | 0.373 | 0.788 | 0.755 | 0.332 | 1.917 |
| **Main Effects** |  |  |  |  |  |  |  |  |  |  |  |  |  |  |
| **Synbiotic** |  |  |  |  |  |  |  |  |  |  |  |  |  |  |
| No |  | 0.21 | 0.36 | 2.79 | 0.50 | 1.16 | 1.12<sup>a</sup> | 0.52 | 2.78 | 0.36 | 0.76 | 0.74 | 0.30 | 1.85 |
| Yes |  | 0.19 | 0.35 | 2.79 | 0.50 | 1.21 | 1.03<sup>b</sup> | 0.52 | 2.74 | 0.37 | 0.79 | 0.76 | 0.33 | 1.92 |
| **Antibiotic** |  |  |  |  |  |  |  |  |  |  |  |  |  |  |
| No |  | 0.20 | 0.35 | 2.79 | 0.50 | 1.16 | 1.12<sup>a</sup> | 0.52 | 2.79 | 0.36 | 0.76 | 0.74 | 0.30 | 1.85 |
| Yes |  | 0.21 | 0.35 | 2.84 | 0.50 | 1.30 | 1.00<sup>b</sup> | 0.52 | 2.80 | 0.40 | 0.84 | 0.76 | 0.32 | 1.99 |
| **Probability** |  |  |  |  |  |  |  |  |  |  |  |  |  |  |
| Synbiotic |  | 0.973 | 0.375 | 0.502 | 0.244 | 0.931 | **0.072** | 0.309 | 0.335 | 0.059 | 0.646 | 0.621 | 0.962 | 0.367 |
| Antibiotic |  | 0.998 | 0.486 | 0.770 | 0.366 | 0.256 | **0.017** | 0.233 | 0.681 | 0.930 | 0.182 | 0.677 | 0.352 | 0.703 |
| Synbiotic × Antibiotic |  | 0.999 | 0.313 | 0.517 | 0.175 | 0.822 | 0.109 | 0.105 | 0.467 | **0.030** | 0.867 | 0.249 | **0.035** | 0.246 |

<sup>1</sup> Pan-Pancreas weight, Pro-Proventriculus, Giz-Gizzard, Duo-Duodenum, Jej-Jejunum, Ile-Ileum, Cae-Caecum, SI-small intestine.
<sup>2</sup>SEM-Standard Error of Mean (n=8).
<sup>a-b</sup>Means within a main effect or interaction not sharing a common superscript are significantly different ( $P \leq 0.05$ ).

As shown in Table 7, a significant synbiotic and antibiotic interaction was observed for duodenal length (P = 0.007), where synbiotic supplementation increased duodenal length in the presence of antibiotics but had no effect in their absence. No other interactions were significant. Among the main effects, synbiotic supplementation reduced ileal content weight (P = 0.033), while antibiotic supplementation also lowered ileal content weight (P = 0.002). In addition, caeca length tended to be reduced by antibiotic supplementation (P = 0.070). The lengths of the jejunum, ileum, and small intestine, as well as digesta weights of the duodenum, jejunum, small intestine, and caeca, were not significantly influenced by either treatment.

**Table 7.** Effects of dietary synbiotics and early-life antibiotics supplementation on digestive tract lengths and content weights (proportional to body weight) of broiler chicken aged 36 days.

| Synbiotic | Antibiotic | Lengths (cm/100g body weight) |  |  |  |  | Content weights |  |  |  |  |
| --- | --- | --- | --- | --- | --- | --- | --- | --- | --- | --- | --- |
|  |  | <sup>1</sup>Duo | Jej | Ileum | SI | Caeca | Duo | Jej | Ileum | SI | Caeca |
| No | No | 1.35<sup>a</sup> | 2.87 | 3.10 | 7.32 | 1.66 | 0.14 | 0.39 | 0.46 | 0.99 | 0.25 |
|  | Yes | 1.26<sup>ab</sup> | 3.07 | 3.13 | 7.46 | 1.52 | 0.10 | 0.49 | 0.25 | 0.85 | 0.20 |
| Yes | No | 1.21<sup>b</sup> | 2.98 | 3.14 | 7.34 | 1.61 | 0.14 | 0.40 | 0.31 | 0.86 | 0.19 |
|  | Yes | 1.38<sup>a</sup> | 3.16 | 3.15 | 7.69 | 1.61 | 0.10 | 0.44 | 0.23 | 0.78 | 0.19 |
| <sup>2</sup>SEM |  | 0.065 | 0.202 | 0.174 | 0.388 | 0.076 | 0.028 | 0.124 | 0.068 | 0.183 | 0.058 |
| **Main Effects** |  |  |  |  |  |  |  |  |  |  |  |
| **Synbiotic** |  |  |  |  |  |  |  |  |  |  |  |
| No |  | 1.31 | 2.97 | 3.11 | 7.39 | 1.59 | 0.12 | 0.44 | 0.36<sup>a</sup> | 0.92 | 0.23 |
| Yes |  | 1.29 | 3.07 | 3.15 | 7.51 | 1.61 | 0.12 | 0.42 | 0.27<sup>b</sup> | 0.82 | 0.19 |
| **Antibiotic** |  |  |  |  |  |  |  |  |  |  |  |
| No |  | 1.28 | 2.93 | 3.12 | 7.33 | 1.63<sup>a</sup> | 0.14 | 0.40 | 0.39<sup>a</sup> | 0.92 | 0.22 |
| Yes |  | 1.32 | 3.12 | 3.14 | 7.58 | 1.56<sup>b</sup> | 0.10 | 0.47 | 0.24<sup>b</sup> | 0.81 | 0.20 |
| **Probability** |  |  |  |  |  |  |  |  |  |  |  |
| Synbiotic |  | 0.034 | 0.583 | 0.828 | 0.976 | 0.509 | 0.872 | 0.928 | 0.033 | 0.479 | 0.294 |
| Antibiotic |  | 0.198 | 0.320 | 0.905 | 0.722 | 0.070 | 0.224 | 0.426 | 0.002 | 0.436 | 0.436 |
| Synbiotic × Antibiotic |  | 0.007 | 0.940 | 0.954 | 0.698 | 0.190 | 0.874 | 0.734 | 0.190 | 0.813 | 0.535 |

<sup>1</sup>Duo-Duodenum; Jej-Jejunum.
<sup>2</sup>SEM-Standard Error of Mean (n=8).
<sup>a-b</sup>Means within a main effect or interaction not sharing a common superscript are significantly different ( $P \leq 0.05$ ).

As shown in Table 8, there were no significant interactions between dietary synbiotics and early-life antibiotic supplementation for spleen weight and bursa weight. Neither spleen weight nor bursa weight was affected by synbiotic or antibiotic treatments.

**Table 8.** Effects of dietary synbiotics and early-life antibiotics supplementation on immune organ weights (proportional to body weight) and hock burns score of broiler chicken aged 35 days.

| Synbiotic | Antibiotic | Spleen weight (%) | Bursa weight (%) |
| --- | --- | --- | --- |
| No | No | 0.09 | 0.16 |
|  | Yes | 0.09 | 0.15 |
| Yes | No | 0.09 | 0.17 |
|  | Yes | 0.10 | 0.17 |
| <sup>1</sup>SEM |  | 0.012 | 0.025 |
| **Main Effects** |  |  |  |
| **Synbiotic** |  |  |  |
| No |  | 0.09 | 0.16 |
| Yes |  | 0.09 | 0.17 |
| **Antibiotic** |  |  |  |
| No |  | 0.09 | 0.16 |
| Yes |  | 0.09 | 0.16 |
| **Probability** |  |  |  |
| Synbiotic |  | 0.815 | 0.650 |
| Antibiotic |  | 0.879 | 0.980 |
| Synbiotic × Antibiotic |  | 0.645 | 0.861 |

<sup>2</sup>SEM-Standard Error of Mean (n=8).

Table 9 presents the effects of dietary synbiotics and early-life antibiotics on the heterophil-to-lymphocyte (H/L) ratio. A tendency for interaction was observed (P = 0.064). In the absence of antibiotic supplementation, synbiotics reduced the H/L ratio compared with the control, whereas under antibiotic supplementation, synbiotics did not alter the H/L ratio.

**Table 9.** Effects of dietary synbiotics and early-life antibiotics supplementation on heterophil-to-lymphocyte ratio of broiler chickens aged 35 days.

| Synbiotic | Antibiotic | Heterophil-to-lymphocyte ratio |
| --- | --- | --- |
| No | No | 1.07<sup>a</sup> |
|  | Yes | 0.43<sup>c</sup> |
| Yes | No | 0.66<sup>b</sup> |
|  | Yes | 0.42<sup>c</sup> |
| <sup>1</sup>SEM |  | 0.111 |
| **Main Effects** |  |  |
| <u>Synbiotic</u> |  |  |
| No |  | 0.75 |
| Yes |  | 0.54 |
| <u>Antibiotic</u> |  |  |
| No |  | 0.87 |
| Yes |  | 0.43 |
| <u>Probability</u> |  |  |
| Synbiotic |  | 0.009 |
| Antibiotic |  | 0.001 |
| Synbiotic × Antibiotic |  | 0.064 |

<sup>2</sup>SEM-Standard Error of Mean (n=8).
<sup>a-c</sup>Means within a main effect or interaction not sharing a common superscript are significantly different ( $P \leq 0.05$ ).

## Discussion

All treatment groups achieved growth rates and feed conversion efficiencies comparable to Cobb 500 performance standards (Cobb-Vantress, 2022), indicating that synbiotic supplementation and early-life antibiotic administration did not compromise production efficiency under experimental conditions. This finding demonstrates that synbiotics, used alone or concurrently with antibiotics, can sustain broiler performance at commercially expected levels.

Body weight gain exhibited variable effects depending on treatment and developmental stage. During the first week, synbiotic supplementation reduced growth, likely reflecting a transient adaptation period in which metabolic resources were redirected toward establishing gut microbial balance rather than tissue deposition (Abd El-Hack et al., 2020). However, from days 7 to 14, a significant synbiotic-antibiotic interaction emerged, with synbiotic-supplemented birds in the absence of antibiotics achieving the highest weight gains, while untreated controls performed the poorest. Synbiotics improve broiler performance through multiple mechanisms, including pathogen suppression, enrichment of beneficial microorganisms, antimicrobial agent production, immune modulation, organic acid production, and reduced gut pH (Abdel-Moneim et al., 2020). Our results suggest that synbiotics exert maximum benefit when administered without concurrent antibiotic exposure, which may otherwise disrupt microbial colonization. Notably, from days 14 to 21, this pattern reversed, with untreated controls achieving the highest gains while synbiotic-only treatment produced the lowest. Despite these fluctuations, cumulative growth to day 35 did not differ significantly among groups, confirming that both interventions ultimately supported comparable overall performance.

Synbiotic supplementation significantly increased feed consumption during days 7 to 14, coinciding with peak weight gain in the synbiotic-only group. Antibiotic supplementation consistently stimulated feed intake across multiple growth phases and for the overall period; however, this did not translate into superior body weight gain, suggesting possible nutrient utilization inefficiencies resulting from alterations in gut microbiota composition (Stanley et al., 2013; Yadav and Jha, 2019). No significant interactions were detected for feed intake, indicating that synbiotic and antibiotic effects on consumption were largely independent.

Feed conversion ratio further clarified these dynamics. Synbiotic supplementation yielded poorer efficiency in week one but improved conversion from days 14 to 21, suggesting that microbial population stabilization enhanced subsequent nutrient assimilation. Previous research employing *Bacillus subtilis*, the probiotic component in the current formulation, has demonstrated improved apparent total tract nutrient digestibility in broilers and laying hens (He et al., 2019; Neijat et al., 2019). In contrast, antibiotic supplementation worsened overall feed conversion ratio, indicating that despite stimulating feed intake, antibiotics did not enhance growth efficiency, possibly due to adverse effects on microbiota composition and nutrient metabolism (Pan and Yu, 2014; Elokil et al., 2020; Greene et al., 2023). These findings suggest that synbiotics may temporarily impair early growth efficiency but ultimately improve feed utilization, whereas antibiotics increase consumption without conferring proportional efficiency gains.

The synbiotic formulation incorporated *Bacillus subtilis* as the probiotic component and *Saccharomyces cerevisiae* as the prebiotic. Previous studies examining these organisms have yielded inconsistent results, with reports of beneficial effects, adverse outcomes, or statistically non-significant impacts on growth performance across different broiler developmental stages (Mutus et al., 2006; Zhang et al., 2005; Gao et al., 2017; Lin et al., 2023). This variability may reflect differences in microbial viability in feed, probiotic load, genetic strain susceptibility, husbandry and environmental conditions, and interactions with other dietary components (Alkhalf et al., 2010; Yang et al., 2012; Jha et al., 2020; Soumeh et al., 2021). Consequently, direct cross-study comparisons present methodological challenges and may not yield reliable conclusions. Research specifically evaluating synbiotic combinations utilizing *Bacillus subtilis* and *Saccharomyces cerevisiae* in broiler nutrition remains limited, and heterogeneity in microbial compositions and experimental protocols further complicates comparative efficacy analyses (Fazelnia et al., 2021).

Gut morphological responses reflect adaptive adjustments to nutrient digestibility. Increased intestinal mass and length generally indicate reduced digestibility, as the gut compensates through tissue expansion and extended digesta retention to enhance nutrient extraction. Conversely, improved digestibility is associated with lighter, shorter intestinal segments reflecting greater absorption efficiency (Brenes et al., 1993; Karunaratne et al., 2021). Synbiotic supplementation increased duodenal length and relative empty weights of the duodenum and caeca exclusively when combined with antibiotics. This interaction suggests that synbiotic-antibiotic co-administration impaired nutrient digestibility, necessitating compensatory enlargement of these segments (Brenes et al., 1993). The duodenum, as the primary enzymatic digestion site, and the caeca, which facilitate microbial fermentation (Elling-Staats et al., 2022; Han et al., 2023), likely underwent hypertrophy to support intensified digestive and absorptive activity under these conditions. Antibiotic supplementation alone reduced ileal weight and tended to shorten caecal length, consistent with suppressed microbial activity and reduced fermentable substrate availability (Kogut, 2019), which diminishes physiological demand for development of these segments.

Digesta content weights provide additional mechanistic insights. Increased digesta mass reflects reduced digestibility, slower transit rates, and microbial and physicochemical processes (Jorgensen et al., 1996; Svihus and Hetland, 2001; Svihus et al., 2002). Active microbial fermentation, particularly when supported by prebiotic substrates, generates biomass and short-chain fatty acids that increase water retention in digesta, thereby elevating content weights (Liu et al., 2021; Song et al., 2022). Yeast cell wall components and oligosaccharides are known to enhance the water-holding capacity of intestinal contents. Microbial metabolites such as lactate and short-chain fatty acids that result from the fermentation of these oligosaccharides may increase osmotic pressure in the lumen (Liu et al., 2021; Xue et al., 2017), drawing water into the gut and further increasing digesta mass (Choct et al., 2010). Conversely, reduced content weights may indicate more efficient nutrient absorption and clearance, faster transit, or diminished microbial activity with less fermentative bulk (Rougière and Carré, 2010). Short-chain fatty acids stimulate the release of peptide tyrosine-tyrosine (PYY), glucagon-like peptide-1 (GLP-1), and glucagon-like peptide-2 (GLP-2) from L-cells in the small intestine (Brooks et al., 2016). Among these, GLP-2 promotes intestinal mucosal hyperplasia (Hu et al., 2010), whereas PYY and GLP-1 activate the ileal brake mechanism, thereby reducing gastric emptying and slowing gastrointestinal motility (Meyer et al., 1998; Maljaars et al., 2008).

Both synbiotics and antibiotics independently reduced ileal content weight through distinct mechanisms. Synbiotics likely improved nutrient digestibility and accelerated ileal digesta clearance by enhancing beneficial microbial populations and activating ileal brake responses via increased short-chain fatty acid production (Mountzouris et al., 2010). Antibiotics reduced microbial biomass and fermentative substrate availability, resulting in decreased digesta content weights (Pedroso et al., 2006; Lin et al., 2013). Antibiotic supplementation also reduced ileal weight and tended to shorten caecal length, consistent with microbial suppression and reduced physiological demand in these regions (Dumonceaux et al., 2006; Gadde et al., 2017).

Overall, synbiotics administered alone did not increase gut weights or lengths, suggesting they did not impair digestibility and may have enhanced efficiency. In contrast, synbiotic-antibiotic combinations induced significant duodenal and caecal enlargement, likely reflecting compensatory responses to reduced digestibility, altered microbial fermentation, and modified osmotic dynamics (Brenes et al., 1993; Choct et al., 2010; Lin et al., 2013).

Neither treatment affected spleen nor bursa weights, suggesting that synbiotics and antibiotics did not markedly influence primary or secondary lymphoid organ development, consistent with studies indicating that functional feed additives often modulate immune responses without necessarily altering gross organ morphology (Madej et al., 2015; Abdel-Hafeez et al., 2017; Yun et al., 2017). Immune function was more clearly reflected in the heterophil-to-lymphocyte ratio, a well-established physiological stress indicator (Thiam et al., 2022; Brathen et al., 2025). Synbiotic supplementation significantly lowered this ratio in the absence of antibiotics, indicating improved stress resilience and immunological stability (Wang et al., 2018; Ding et al., 2019; Song et al., 2022). This effect likely reflects enhanced gut microbiota balance, increased production of immunomodulatory metabolites such as short-chain fatty acids and reduced systemic stress signalling via the microbiota-gut-brain axis (Villageliu and Lyte, 2017; Cao et al., 2021; Liu et al., 2021). When antibiotics were co-administered, this synbiotic benefit disappeared, suggesting that microbial suppression impaired the ability of synbiotics to promote immune homeostasis (Costa et al., 2017; Elokil et al., 2020). Antibiotic supplementation alone also reduced the heterophil-to-lymphocyte ratio; however, this likely reflects pharmacological suppression of microbial antigenic stimulation rather than true enhancement of immune resilience (Kogut, 2019; Yadav and Jha, 2019). These results suggest that synbiotics function as effective immunomodulators in antibiotic-free environments, whereas concurrent antibiotic administration masks or blunts their beneficial effects.

Synbiotics sustained Cobb 500 performance standards across all treatments (Cobb-Vantress, 2022) but demonstrated distinct advantages only in the absence of antibiotics. Under antibiotic-free conditions, synbiotics improved mid-growth feed efficiency and reduced physiological stress, evidenced by lower heterophil-to-lymphocyte ratios, without inducing compensatory gut hypertrophy. These results indicate that synbiotics promoted more efficient nutrient utilization and enhanced immune homeostasis when the gut microbiota established naturally. Conversely, synbiotic-antibiotic combinations induced increased duodenal and caecal development, indicative of compensatory responses to impaired digestibility, without improving overall performance or immune balance. These findings suggest that antibiotics undermine the functional efficacy of synbiotics by altering microbial colonization and shifting host resource allocation toward maintaining gut function at the expense of systemic resilience.

## Conclusions

Synbiotics administered without antibiotics improved mid-growth feed efficiency and reduced physiological stress, establishing their value as functional alternatives in antibiotic-free production systems. However, concurrent antibiotic use induced compensatory gut hypertrophy without improving overall performance or immune balance, suggesting that antibiotic co-administration compromises synbiotic efficacy. These findings support the targeted application of synbiotics as sustainable antibiotic replacements in broiler production while emphasizing the need for refined management practices to optimize production, health, and welfare outcomes. Further research examining nutrient digestibility, fermentation dynamics, and gut microbiota composition is necessary to elucidate the mechanistic pathways underlying the interactive effects of antibiotics and synbiotics on broiler performance and welfare.

## Acknowledgements

The financial assistance for the research project was provided by Bio Nutri International (Pvt) Ltd. Acknowledgment is granted to the technical staff of the Department of Farm Animal Production and Health, Faculty of Veterinary Medicine and Animal Science, and Department of Animal Science, Faculty of Agriculture at the University of Peradeniya for their assistance in conducting the animal trial and laboratory analysis. The support given by the slaughterhouse staff of the Udaperadeniya Livestock Farm, Faculty of Agriculture of the University of Peradeniya, is also acknowledged.

